# Burden of rare pathogenic variants suggests disrupted cytoskeletal organisation in the pathogenesis of pulmonary fibrosis

**DOI:** 10.1101/2024.06.10.598229

**Authors:** Konain Bhatti, Dapeng Wang, Hernan Fainberg, Andressa Alves Bordignon, Yujie Ni, Bin Liu, Alison John, Richard Allen, Louise V. Wain, Simon R. Johnson, Toby M Maher, Philip L Molyneaux, Elisabetta Renzoni, Gauri Saini, Deborah Morris-Rosendahl, R Gisli Jenkins, Iain Stewart

## Abstract

**Background:** Rare genetic variants contribute to pulmonary fibrosis (PF) risk and outcome, with known variants highlighting the importance of impaired telomere maintenance and surfactant biology. However, much of the disrupted genetic architecture of PF remains unexplained. This study aimed to identify genes with rare pathogenic coding variants that represented a burden at the exon level associated with the pathogenesis of PF.

**Methods:** PF case cohorts included the PROFILE study of incident idiopathic pulmonary fibrosis (IPF) and PF-classified participants from the Genomics England 100K (GE100KGP) study. Whole genome sequencing data were used to test the burden of rare protein altering variants (PAVs; CADD score >20) defined by minor allele frequency <0.1% in the summary level gnomAD reference database (v3.1.2). A summary exon-level pathogenicity score was derived by standardising PAV predictions from functional annotation tools (AlphaMissense, REVEL, ClinPred, CADD, PolyPhen-2, SIFT) averaged across exons. SKAT-O based kernel regression associated exon-level pathogenicity features with disease progression and survival. Single cell lung transcriptomic datasets were harmonised to evaluate expression patterns and gene ontology characterised enriched molecular functions.

**Results:** Across 507 PROFILE cases and 451 GE100KGP cases, each compared against 76,156 reference participants, 77 genes showed overlapping significant rare PAV burden, comprising 206 concordant PAVs. Of these, 15 genes had exons with pathogenicity scores that were associated with worse clinical outcomes in IPF, including *FAT4* exon-10 and *DNAH7* exon-43. Dysregulated gene expression of *COL6A3* and *FAT4* was observed in IPF lung fibroblasts, while *DNAH7*, *DNAH12* and *PCM1* showed high expression in IPF epithelial cells. The top enriched molecular functions were related to cytoskeletal motor activity.

**Conclusions:** Rare PAV burden highlighted that pathogenicity within specific exons was associated with PF risk and worse clinical outcomes, identifying genes with dysregulated expression in disease lung cells. The genetic architecture implicates disrupted cytoskeletal organisation in the pathogenesis of PF.

## Introduction

Pulmonary fibrosis (PF) is a chronic, progressive process characteristic of many interstitial lung diseases (ILD) including idiopathic pulmonary fibrosis (IPF) and familial pulmonary fibrosis (FPF)[1]. Although antifibrotic treatments such as pirfenidone and nintedanib can slow disease progression, people with PF continue to experience poor prognosis and limited survival after diagnosis[2]. The considerable overlap in clinical characteristics, genetic variants and environmental risk factors between IPF and FPF[3–5], suggests common molecular mechanisms underlie PF pathogenesis.

Genome-wide association studies (GWAS) have identified common variants in coding and non-coding regulatory regions linked to IPF susceptibility, with over 30 signals associated with increased risk[6]. Additional analyses have identified common variants associated with prognosis, including *PCSK6* which was associated with IPF survival[7], and *PKN2* with longitudinal lung function decline of IPF[8].

Whole genome sequencing (WGS) supports identification of rare pathogenic variants across the genome, including coding regions in which variants may lead to truncation and loss of function that alters protein function as well as changes in gene expression. Whole exome sequencing (WES) has highlighted rare missense variants in genes implicating mitotic checkpoint signalling as a dysregulated pathway in fibrotic development[9, 10], whilst rare variants in surfactant genes identified in sporadic PF have also been identified in FPF[11]. Telomere-related rare variants have been associated with worse survival in two independent IPF cohorts, increasing estimated risk of mortality by 57% in meta-analysis[12], and explain up to 23% of FPF risk[13]. Despite advances in understanding the genetic architecture of PF, a large proportion of genetic risk and genetically dysregulated molecular mechanisms are yet to be elucidated.

This study aims to prioritise biologically plausible genes and pathways that contribute to PF susceptibility and progression by evaluating the burden of rare pathogenic protein-altering variants (PAVs), consolidating the predicted pathogenicity at exon-level, and interpreting these findings in the context of transcriptomic signals and molecular pathways.

## Methods

### Populations

WGS data were obtained from the PROFILE cohort, a UK based prospective study of incident IPF diagnoses[14], and the Genomics England 100K Genome Project (GE100KGP), a UK initiative aiming to produce approximately 100,000 human genomes using WGS to study the causes of cancer and rare diseases. Analysis of GE100KGP data was performed within the Research Environment of Genomics England. The case samples in GE100KGP cohort were defined based on data release v14 as those recruited to the rare disease FPF programme or patients with Human Phenotype Ontology (HPO) codes HP:0002206, HP:0006530, or International Classification of Diseases (ICD-10) codes J84.1, J84.9, J84.8, J67.9 (Supplementary Table S1). Quality control procedures applied to PROFILE and GE100KGP case cohorts included sex concordance, confirmation of European ancestry and exclusion of related individuals[9] (Supplementary Methods). Control data were derived from summary level variant call format (VCF) files from the Genome Aggregation Database (gnomAD v3.1.2)[15], comprising 76,156 individuals with WGS aligned to the GRCh38 reference genome without enrichment of any specific diseases or restriction on ancestry.

### Rare variant burden testing

Rare variants were defined as variants with an allele frequency < 0.1%. Additional filtering for internal allele frequency cut-off was also applied (Supplementary Table S2). Variants in coding regions were defined as protein-altering or synonymous based on the consequence predicted by the Ensembl Variant Effect Predictor (VEP) and Combined Annotation–Dependent Depletion (CADD) score provided within the gnomAD VCF files[16]. The Testing Rare vAriants using Public Data (TRAPD) pipeline was used for the burden testing between the individual level case data and summary level control data[17]. Potentially pathogenic rare variants were identified, the relationships between the variants and genes were constructed, and the number of individuals who carried the specific alleles was calculated for both case and control cohorts. Burden testing was performed by using the Fisher exact test at P-value < 1 × 10^-10^ under the dominant model. Genes that were significant for synonymous variants were excluded from subsequent prioritisation analysis to reduce false discovery (Supplementary Table S2).

### Exon-level burden assessment

Genes showing significant rare PAV burden in both cohorts relative to the gnomAD reference in TRAPD analyses were taken forward. Variants were filtered using gnomAD sequencing coverage metrics, excluding loci with insufficient coverage (depth ≥20× in <50% of gnomAD samples). To support exon-level aggregation, genes were retained only if they contained at least one QC-passing rare PAV within the same exon in both cohorts. Predicted pathogenicity for individual rare variants was summarised using established in-silico predictors, Polyphen-2[18], SIFT[19], CADD[20], alphaMissense[21], ClinPred[22], REVEL[23]. Outputs from each tool were normalised to a common scale 0-1 (higher values indicating greater predicted pathogenicity) and summarised as a variant-level pathogenicity score (range 0-6). Agreement between the summary score and predictor outputs was assessed with Pearson correlation, r=0.7 with AlphaMissense, REVEL, ClinPred and PolyPhen-2 (Supplementary Figure S1). Exon-level burden was defined as the mean summary pathogenicity score of rare PAVs within each exon, with exons retained in downstream analysis if mean score ≥1.5 and containing at least one concordant rare PAV across case cohorts.

### Functional association of exon-burden with prognosis

The PROFILE prospective observational cohort of incident IPF participants includes disease progression, defined as 10% relative decline of percent predicted forced vital capacity (ppFVC) or mortality within 12 months, and overall survival. In downstream exon-level association analyses, rare PAVs within retained exons were collapsed into a per-participant exon burden score summarising the cumulative predicted pathogenicity of carried variants. SKAT-O was used as a kernel regression framework applied to exon burden features to evaluate whether variation in exon burden scores was associated with prognosis. The individual-by-exon burden matrix was used as the input feature matrix for SKAT-O to test exon-level associations with dichotomous endpoints of disease progression and survival below median, adjusting for age. Region-based associations have been similarly applied in other diseases[24]. Standardised effect sizes were derived using the squared Wald statistic (F-statistic), and hierarchical clustering was performed using Ward linkage on Euclidean distance (SciPy) to visualise functional relationships. An f-statistic ≥1.5 was interpreted to represent a model in which the variance explained exceeded the residual error by at least 50% and used to select associations. Disease outcomes were unavailable in GE100KGP.

### Differential gene expression and pathway analyses

Three publicly available single cell RNA sequencing (scRNA-seq) lung datasets were retrieved and integrated to create a representative dataset; GSE136831[25], GSE135893[26], and GSE128033[27] were obtained from the Gene Expression (GEO) Omnibus (GEO). To support consistency with the PROFILE cases in functional associations, only IPF and control lung samples were included (Supplementary Methods). Dataset integration was performed using Harmony to align multi-dataset cells while preserving biologically meaningful variability (Supplementary Figure S2-4)[28]. scRNA-seq preprocessing, clustering, and data handling were performed in Seurat, with cell-type annotation guided by the Human Lung Cell Atlas. Differential gene expression analysis was performed with DESeq2 pseudobulk aggregation per donor per cell type. To compare expression of genes across cell types and disease status, z-scores were calculated from average expression values of genes within each annotated cell type regardless of disease status and plotted for IPF samples. Gene Ontology (GO) molecular function term enrichment was assessed using g:Profiler, the top ten significant terms were plotted using -log10 transformed p-values.

## Results

### Pathogenic rare variant burden by case

After pre-processing of 541 WGS PROFILE samples, 507 case samples were used in downstream analysis. A total of 451 PF case samples from GE100KGP were used in downstream analysis. The average age of people in PROFILE cases was 70.5 years (±8.4), which was older than the GE100KGP cases of 65.5 years (±14.5), whilst PROFILE cases were also represented by more males than GE100KGP cases, 76.3% vs 51.4%, respectively (Supplementary Table S3).

In the PROFILE cohort, 371 genes contained PAVs and were significantly associated with PF (Figure 1A). In the GE100KGP cases, 361 genes contained PAVs and were significantly associated with PF (Figure 1B). Initially, there were 87 genes that were overlapping across both case cohorts (Figure 1C), which were reduced to 77 genes meeting coverage criteria. Overlapping genes were represented by 997 rare PAVs in PROFILE and 920 in GE100KGP, including 206 variants that were concordant across both cohorts (Figure 1D).

**Figure 1.**
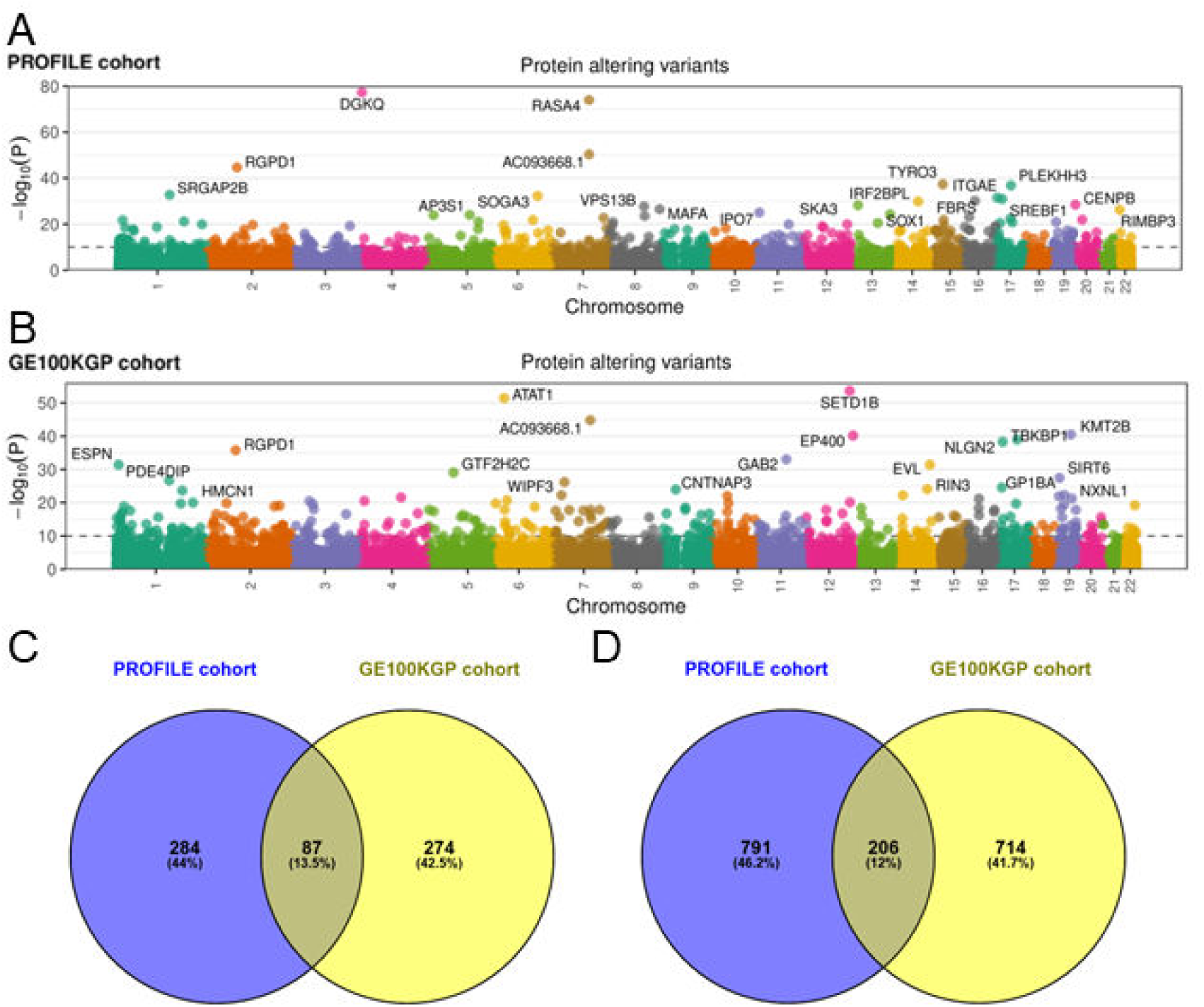
Overlapping rare variant burden of pulmonary fibrosis susceptibility. Coding regions using whole genome sequencing in TRAPD pipeline and gnomAD control in A) PROFILE IPF cohort B) Genomics England 100K Genome Project (GE100KGP) PF cohort. C) Venn diagram of overlapping genes with significant rare protein altering variant burden between A and B. D) Venn diagram of overlapping rare protein altering variants with sufficient gnomAD coverage; represents 77/87 shared genes in C.

A significant burden of protein-altering variants was observed in both PROFILE and GE100KGP cohorts for the known telomere-associated risk genes *TERT* (p=7.87E-15, p=1.36E-18, respectively) and *RTEL1* (p=4.08E-17, p=8.42E-15, respectively) (Supplementary Table S4). Following exclusion based on gnomAD coverage, there were 5 unique *RTEL1* variants in PROFILE (case n=6) and 3 in GE100KGP (case n=3). *TERT* was retained in PROFILE (2 unique variants, case n=2), but not GE100KGP.

### Exon-level summary burden

There were 40 genes that were represented by 68 exons with mean summary exon-level pathogenicity ≥1.5 and at least one concordant PAV between case cohorts. In PROFILE, 168 variants were retained and in GE100KGP, 162 variants were retained. The greatest mean summary pathogenicity of 4.39 was observed for *DNAH7* exon 43 (Figure 2A). Twenty-two exons (32%) had a mean pathogenicity ≥3. Twelve gene exons included >1 concordant variant across cohorts, *FAT4* exon-2 included four concordant variants (mean pathogenicity 2.32). In the PROFILE cohort, the gene exons with the most frequent case burden of variants were *KIF26A* exon-12 (case n=15), followed by FAT3 exon-10 (case n=12), and *RANBP2* exon-20 (case n=10) (Figure 2B). In the GE100KGP cohort, *KIF26A* exon-12 variants and *FAT3* exon-10 variants were similarly represented by the most frequent number of cases (9 and 8, respectively), whilst *MYO15A* exon-2 variants were also observed in 8 cases (Figure 2B).

**Figure 2.**
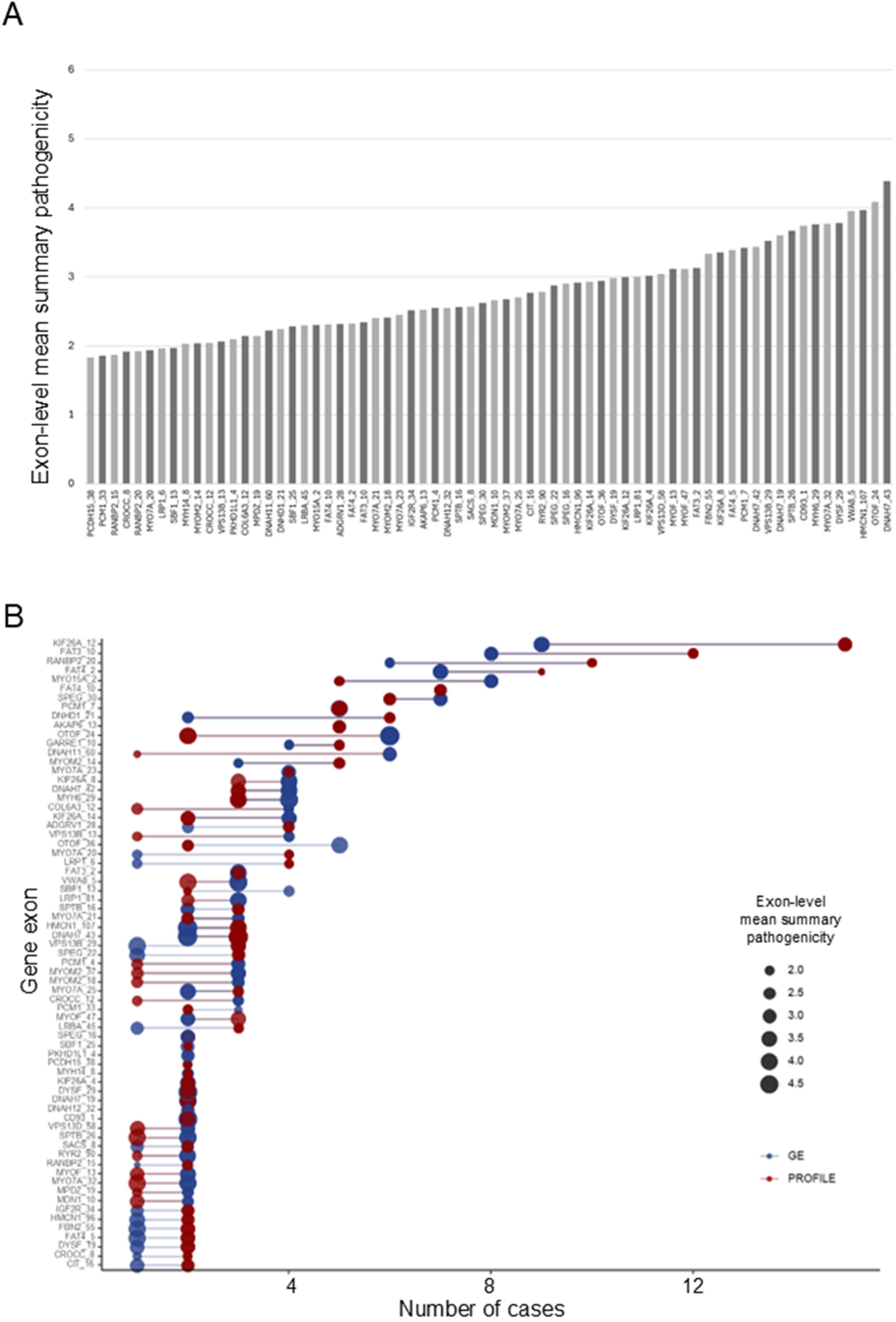
Exon-level pathogenicity and case numbers across case cohorts. A) Exon mean summary variant pathogenicity score presented for 68 gene exons. B) Exon mean summary variant pathogenicity score and case number represented in PROFILE and GE100KGP for 68 gene exons.

### Association between exon-level burden and prognosis

Outcome modelling in the PROFILE cohort was restricted to carriers of qualifying variants within the 68 exons, representing 164 participants, of which 75 participants (45.7%) exhibited disease progression within 12 months and for whom median survival was 3.9 years (95%CI 2.8 to 4.8). Seventeen exons, representing 15 genes, clustered together in weighted association with greater risk of worse clinical outcome (Figure 3, Table 1, Supplementary Figure S5). Three exons, in three distinct genes, were associated with both disease progression and survival, including *FAT4* exon-10, *DYSF* exon-29, and *MYH6* exon-29. *DYSF* exon-19 was primarily associated with disease progression. An association was observed between *KIF26A* and both disease progression and survival endpoints, with exon-14 being associated with disease progression and exon-12 associated with survival. Strong association was suggested between *DNAH7* exon-43 and disease progression alone, whilst associations were also observed with *PCDH15* exon-38, *DYSF* exon-19, *SBF1* exon-13, *PCM1* exon-4, *MYOM2* exon-37, *VPS13D* exon-58, *VPS13B* exon-13, and *COL6A3* exon-12. *DNAH12* exon-32 was associated with survival alone, associations were also observed with *MYO7A* exon-25 and *MYOF* exon-47.

**Figure 3.**
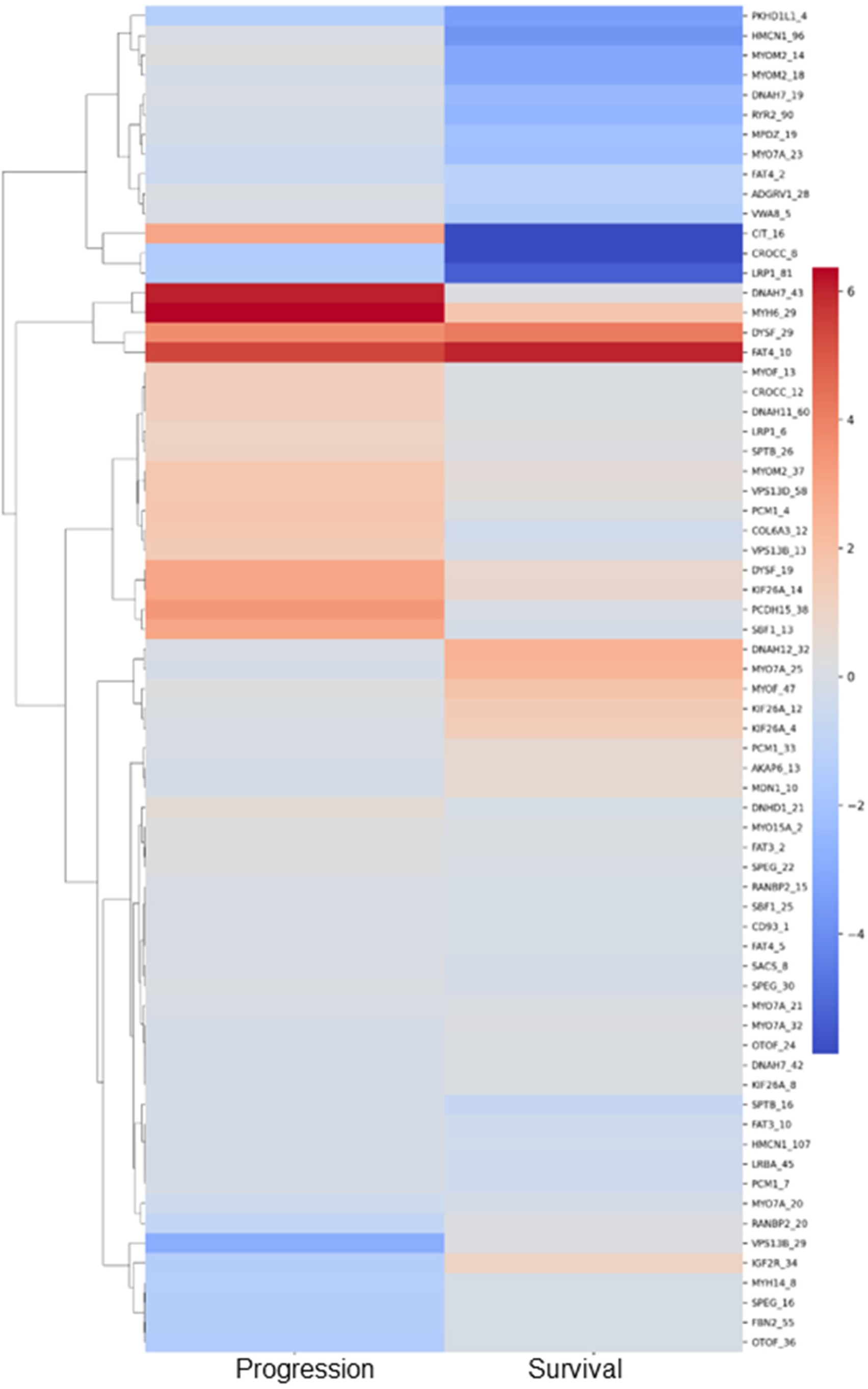
Exon-level pathogenicity weighted association with prognosis in IPF. Heatmap of standardised effect size of exon mean summary pathogenicity (red greater risk of outcome, blue lower risk of outcome) on outcomes of disease progression (prog: 10% relative decline in ppFVC or mortality within 12 months) or survival below median (surv: median survival 3.9 years) in PROFILE cases (n=164) derived from SKAT-O based kernel regression. 17/68 gene exons, representing 15 genes, met F-statistic ≥1.5 for risk of at least one outcome without a protective effect on other.

**Table 1:**
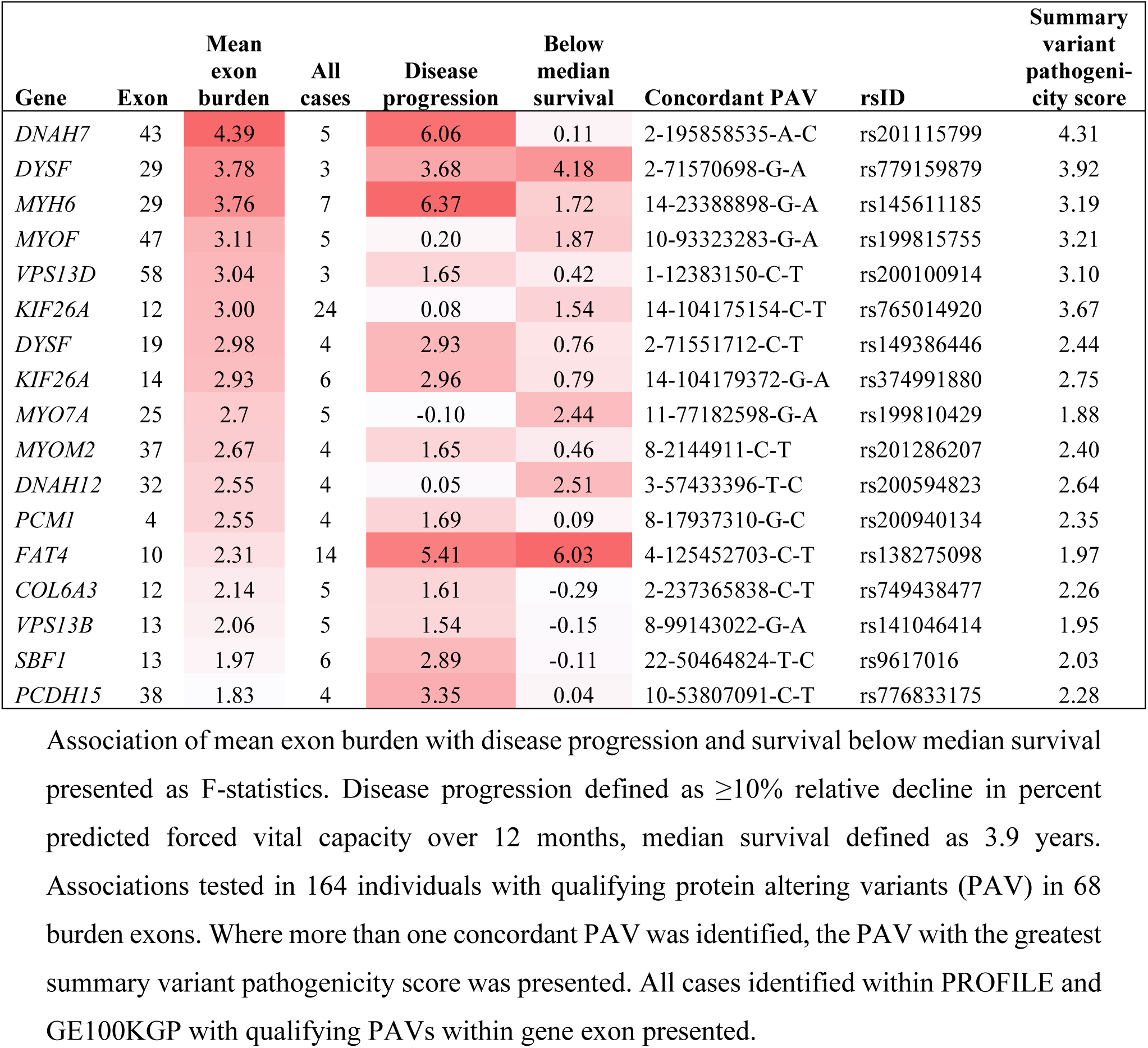
Prioritised burden genes involved in PF pathogenesis.

### Dysregulated expression and enriched pathways of burden genes

Differential gene expression across lung cell types supported dysregulation of the 15 exon-level burden genes in IPF lung tissue. *COL6A3* and *FAT4* were upregulated in IPF stromal lineages (Supplementary Figure S6), including fibroblasts and myofibroblasts (Figure 4A). *DNAH7*, *DNAH12*, *PCM1* were primarily upregulated in IPF epithelial lineages (Supplementary Figure S7), particularly multiciliated cells, whilst *MYOF* was upregulated in AT1 cells (Figure 4A). *KIF26A*, *DYSF*, *PCDH15*, as well as *FAT4*, were upregulated in IPF endothelial cells, particularly lymphatic differentiating endothelial cells (Figure 4A, Supplementary Figure S8). *SBF1* was upregulated in IPF lymphatic proliferating endothelial cells; *VPS13B*, *VPS13D* and *MYOM2* were upregulated in IPF ‘SM activated stress response’ cells (Figure 4A, Supplementary Figure S9). *MYO7A* and *MYH6* demonstrated the lowest expression across lung cells (Supplementary Figure S10).

**Figure 4.**
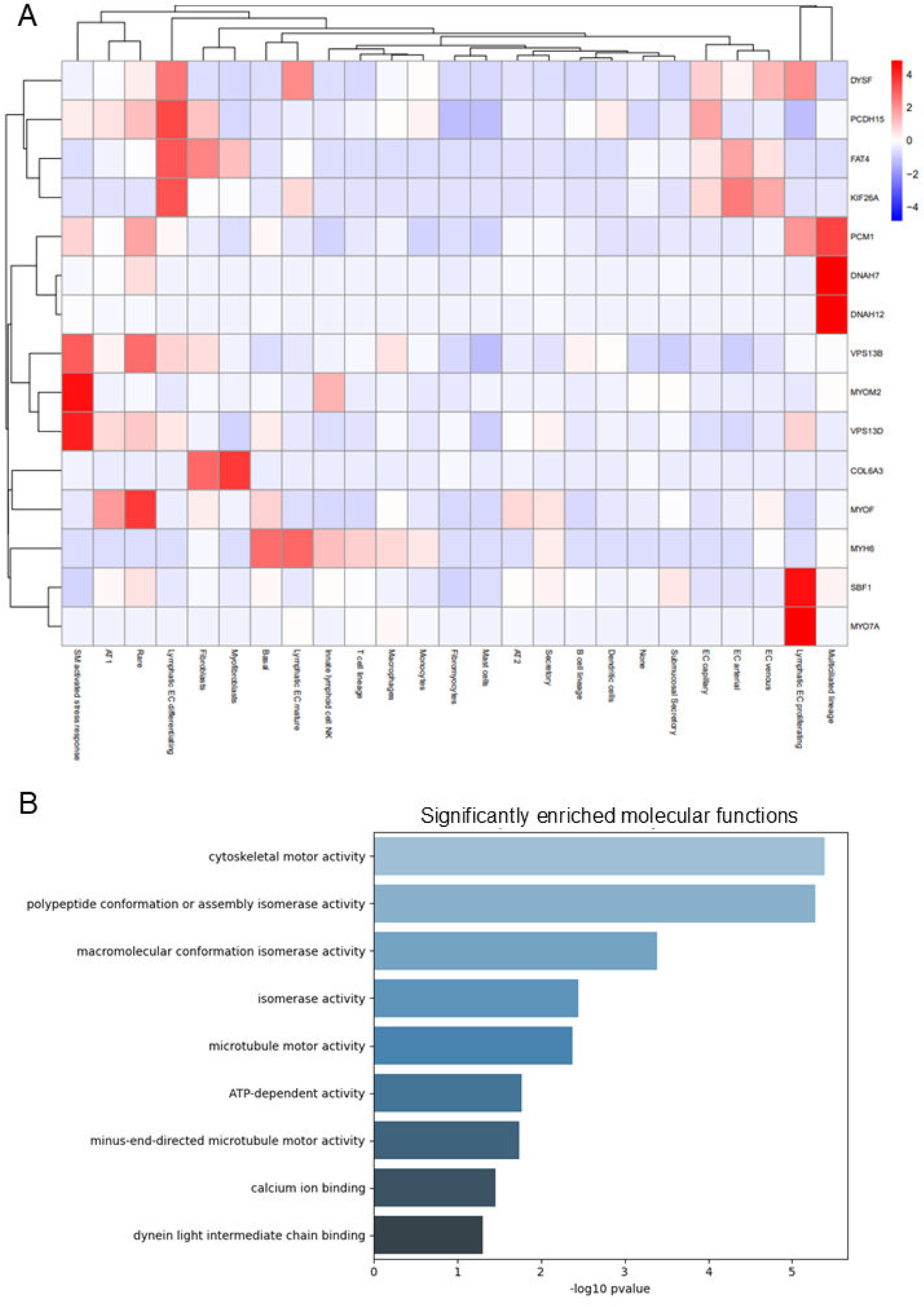
IPF lung cell expression and enriched functions of burden genes. A) Heatmap of differential gene expression IPF z-scores in integrated single cell RNA sequencing for 15 genes with F-statistics ≥1.5, annotated at Human Lung Cell Atlas level 3. Red represents strongly upregulated relative to respective gene average, blue represents strongly downregulated. B) Enrichment analysis for 15 genes, Gene Ontology presents significant molecular function terms.

Enrichment analysis performed on 40 genes with exon-level burden identified molecular function of cytoskeletal motor activity (p=4.63E-10) and polypeptide conformation or assembly activity (p=7.38E-10) within the top pathways, as well as related functions such as microtubule motor activity (p=9.70E-05) and dynein binding, which was consistent with 77 overlapping PAV burden genes (Supplementary Figure S11). The top enriched molecular functions for the 15 genes with exons associated with worse prognosis continued to be cytoskeletal motor activity (p=4.05E-06), polypeptide conformation and assembly activity (p=5.23E-06), and other functions including microtubule motor activity (Figure 4B).

## Discussion

Approximately 75% of FPF cases have no known genetic cause, whilst 10% of sporadic PF cases are thought to be hereditary, suggesting a spectrum of genetic risk with substantial unexplained genetic architecture and pathogenesis[13, 29]. By focusing on reproducible rare PAV burden in susceptibility across two case cohorts and consolidating variant effects at the exon level that were associated with poorer outcomes, we prioritised 15 genes. These genes were consistently dysregulated in IPF lung single-cell datasets and converged on cytoskeletal motor and microtubule-related molecular functions.

We identified that *COL6A3* and *FAT4* were genes with exon-level burden of pathogenic rare variants associated with poor outcomes along with dysregulated gene expression in stromal lineages of fibroblasts and myofibroblasts. *COL6A3* encodes collagen-alpha-1(VI), an extracellular matrix (ECM) component and an established biomarker in IPF[30, 31]. Greater levels of collagen in the IPF ECM leads to increased stiffness, whilst mechanical strain releases ECM-sequestered profibrotic cytokine TGFβ. *FAT4* encodes an atypical cadherin protein that is part of the canonical Hippo signalling pathway involved in proliferation, differentiation, and increasingly implicated in IPF pathogenesis[32, 33]. FAT4 is localised to primary cilia, a signalling structure distinct from motile cilia that hold critical functions in orientation of cell division and fibroblast migration during wound healing [34, 35], and mediate responses to ECM perturbations[36]. Compared with normal lung, greater numbers of primary cilia have been observed in IPF lung fibroblasts, which are also characterised by short primary cilia morphology that can be induced by TGFβ and restored by microtubule stabilisation[37, 38]. The rare variant observations support a genetic architecture in which the interactions between fibroblast organisation and ECM properties are dysregulated.

Microtubule organisation components showed increased gene expression in IPF multi-ciliated epithelial cells. Dynein axonemal heavy chain-7 and -12, encoded by *DNAH7*, *DNAH12* respectively, are microtubule-associated motor proteins that control cilium movement, whilst microtubule anchoring protein pericentriolar material-1, encoded by *PCM1*, is involved in recruiting proteins to centriolar satellites, influencing microtubule organisation, ciliogenesis and cilia motility[39]. Mutations in outer dynein arms have been associated with primary ciliary dyskinesia, a rare recessive disorder in which defective motile cilia result in poor mucociliary clearance, leading to chronic respiratory problems including persistent infection, inflammation, and increasing bronchiectasis with age. As PF is frequently diagnosed in late adulthood, the penetrance of rare variants may be incomplete without accumulation of other lifetime exposures[40]. Mucus hypersecretion and ciliary impairment in airways have been linked to the development of mucus plugs in alveolar regions of IPF patients[41], whilst lung fibrosis development in a bleomycin model was reduced by modulating cilia structure in Krt5+ basal cells[42]. The genetic architecture of microtubule disorganisation in multiciliated cells may predispose individuals to impaired mucociliary transport, chronic proinflammatory signalling and recurrent microinjury, manifesting fibrotic remodelling over the life course of an individual. With established evidence that the *MUC5B* common variant is the strongest associated genetic risk factor for IPF susceptibility[6], this rare variant burden suggests airway mucociliary function may be more important in fibrogenesis than currently attributed.

Dysregulated expression of burden genes *KIF26A*, *DYSF*, *PCDH15*, *FAT4*, and *SBF1*, was observed in lymphatic endothelial cells. Studies have shown that endothelial cells can transition and contribute to the myofibroblast pool in pulmonary fibrosis models, corroborated through cell fate mapping in human IPF scRNAseq[43]. Lymphangiogenesis has been implicated in fibrotic lung remodelling, with the diameter of lymphatic vessels in alveolar spaces of IPF lung tissue correlating with disease severity[44, 45]. Protocadherin-15, kinesin family member 26A, and SET binding factor-1, encoded by *PCDH15*, *KIF26A* and *SBF1* respectively, as well as FAT4 described above, have roles in cell adhesion, cytoskeletal organisation and cellular morphology that could indicate a role in endothelial and lymphatic remodelling[46]. *DYSF,* encoding dysferlin, is an essential regulator of vascular homeostasis through its control of endothelial cell adhesion and membrane integrity [47], highlighting vascular remodelling in the genetic architecture of PF.

Myoferlin, encoded by burden gene *MYOF* and member of the Ferlin family with homology to dysferlin, was dysregulated in IPF AT1 cells. In IPF, AT2 to AT1 transition is impaired during alveolar epithelial repair, whilst epithelial metaplasia can lead to the recruitment of myofibroblasts and formation of fibrotic foci[48]. Myoferlin has been shown to induce the transition of epithelial cells into myofibroblasts through control of TGFβ signalling[49], and has previously featured in the top 20 genes during exploratory rare variant burden testing in an independent FPF cohort, although enrichment did not reach statistical significance[13].

Rare variant burden aggregated at exon-level has identified genes involved in cytoskeletal organisation and lung tissue remodelling, including regulation of fibroblastic properties, ECM properties, and mucociliary clearance. This study used variant-level, exon-level, gene-level and pathway-level approaches to iteratively prioritise biologically plausible mechanisms of perturbation within the genetic architecture of PF. However, the approach does not identify rare variant burden independently related to clinical outcomes and, whilst gene expression was evaluated, it is possible that identified PAVs do not alter transcript levels.

Two large PF cohorts were included to support reproducible risk susceptibility variants and exon burden. Inclusion of both familial and sporadic disease can provide insights into converging pathogenic mechanisms. Identification of qualifying variants in *TERT* and *RTEL1* was consistent with previous studies although genomic coverage in gnomAD could not confirm *TERT* variants observed in GE100KGP. No restriction on ancestry was applied to summary level reference data from gnomAD, which may impact rare variant burden in susceptibility, however, associations with disease outcomes were performed within cases of European ancestry with adjustment for age thereby minimising the effect of population stratification on gene prioritisation. No exclusion criteria were applied to cases based on carriers of known causal variants, potentially reducing power to identify novel variants. Reducing technical artifacts and neutral background by excluding genes with significant synonymous burden may have limited identification of biologically relevant signals in very large genes. The findings require functional in-vitro and in-vivo studies to validate the role of genes, variants and their clinical relevance.

## Conclusion

Through selection of reproducible PAV burden in coding regions across two large PF case cohorts, aggregating burden over exons and modelling associations between pathogenicity and disease outcomes, this study identified 15 genes implicated in the genetic architecture of PF susceptibility and prognosis. Using integrated IPF lung transcriptomics, cell-type specific dysregulation was observed in all 15 genes, particularly multiciliated epithelial cells, fibroblasts, and lymphatic endothelial cells, whilst molecular functions relating to cytoskeletal dynamics were enriched. Preclinical functional genomic studies should establish the role of the prioritised genes in the pathogenesis of PF.

## Acknowledgements

This work was funded by an MRC Programme Grant (MR/V00235X/1) awarded to RGJ and a NIHR Imperial Biomedical Research Centre Springboard Award (PSR547) obtained by IS. DW is supported by the Taishan Scholars Program of Shandong Province (tsqn202312110) and the Fundamental Research Funds for the Central Universities (25CX06001A). This research is partly funded by the National Institute for Health and Care Research (NIHR) and Leicester (BRC). The views expressed are those of the author(s) and not necessarily those of the NIHR or the Department of Health and Social Care. This research used the ALICE High Performance Computing Facility at the University of Leicester. This research was made possible through access to data in the National Genomic Research Library, which is managed by Genomics England Limited (a wholly owned company of the Department of Health and Social Care). The National Genomic Research Library holds data provided by patients and collected by the NHS as part of their care and data collected as part of their participation in research. The National Genomic Research Library is funded by the National Institute for Health Research and NHS England. The Wellcome Trust, Cancer Research UK and the Medical Research Council have also funded research infrastructure.

## Contributions

DW, RGJ and IS conceived and designed the study. KB, DW, HF, AAB, YN, IS performed the data analysis. RGJ and IS secured funding. SRJ, TMM, PLM, GS contributed to the patient recruitment and data collection. KB, DW and IS wrote the initial draft. BL, AJ, HF, YN, AAB, LVW, DMR, RGJ, TMM, PLM, EAR, GS and provided input and editing.

