## Supplementary Material for "Burden of rare pathogenic variants suggests disrupted cytoskeletal organisation in the pathogenesis of pulmonary fibrosis"

### Supplementary methods

#### Genome preprocessing

In GE100KGP, samples with concordance between phenotypic sex and inferred ploidy, unrelated individuals, and European ancestry (probability > 0.8) available in GRCh38 reference genome were retained. BCFtools was used for the processing of VCF files into a single VCF file per cohort, keeping only variants that passed quality controls. Multi-allelic variants were split and indels were left-aligned and normalised. The missing genotypes were replaced with reference alleles. For both case and control samples, variants from 22 autosomes were selected and unique variant identifier was constructed to link the variants across cohorts. To support discovery analysis using burden testing, variants that were shared between case and controls and case specific variants were included.

#### Transcriptomic preprocessing

Established processing steps were followed for each dataset, including obtaining the number of genes detected per UMI and the mitochondrial ratio. For GSE136831 and GSE128033 datasets, cell-level filtering included removing cells with mitochondrial ratios less than 20% and maintained cells with >500 UMIs per cells and >1000 genes per cell. For GSE135893 dataset, a different set of thresholds were used: mitochondrial ratio <15%, UMIs >1000 and >1000 genes per cell. Gene-level filtering for all datasets included removing genes with zero reads for all cells and keeping genes with expression in ≥10 cells. Log-normalisation was performed on each dataset, followed by variance stabilisation using SCTransform. To remove the batch effect arising from the inter-sample variations, the integration analysis was carried out by finding 2000 highly variable genes and employing the mutual nearest neighbours method [1]. Harmony was used to integrate the multi-dataset cells into clusters [2], scRNA-seq preprocessing, clustering, and data handling were performed in Seurat [3]. Cell-type annotation was performed with the Human Lung Cell Atlas [4], gene expression analysis was performed with DESeq2 [5].

### Table S1. Definition of PF case group for GE100KGP cohort.

| System | Disease term | Code | Number of samples (N=451) |
|---|---|---|---|
| Rare Disease | Familiar pulmonary fibrosis | | 111 |
| HPO | Pulmonary fibrosis | HP:0002206 | 105 |
| HPO | Interstitial pulmonary abnormality | HP:0006530 | 81 |
| ICD-10 | Other interstitial pulmonary diseases with fibrosis | J84.1 | 344 |
| ICD-10 | Interstitial pulmonary disease, unspecified | J84.9 | 135 |
| ICD-10 | Other specified interstitial pulmonary diseases | J84.8 | 30 |
| ICD-10 | Hypersensitivity pneumonitis due to unspecified organic dust | J67.9 | 14 |

### Table S2. Models for burden testing.

| Model | External AF cut-off | Internal AF cut-off | CADD score | Variant type | Variant inclusion |
|---|---|---|---|---|---|
| Rare Protein Altering Variants | AF < 0.1% | AF < 1% | >20 | PAV | splice_acceptor_variant; splice_donor_variant; stop_gained; frameshift_variant; stop_lost; start_lost; inframe_insertion; inframe_deletion; missense_variant; protein_altering_variant |
| Rare synonymous | AF < 0.1% | AF < 1% | <=20 | synonymous | synonymous_variant |

### Table S3. Case demographics

| System | PROFILE | GE100KGP |
|---|---|---|
| Total cases in analysis | 507 | 451 |
| Mean age (SD) | 70.5 (8.4) | 65.5 (14.5) |
| Male (%) | 387 (76.3%) | 232 (51.4%) |

**Table S4. 77 genes with overlapping rare variant burden between case cohorts**

| Gene name | Gene ID | PROFILE p-val | GE100KGP p-val |
|---|---|---|---|
| *ADAMTSL1* | ENSG00000178031 | 4.01E-11 | 2.51E-12 |
| *ADAMTSL2* | ENSG00000197859 | 2.18E-11 | 4.32E-12 |
| *ADGRV1* | ENSG00000164199 | 1.00E-15 | 1.91E-16 |
| *AGAP3* | ENSG00000133612 | 6.26E-14 | 5.53E-15 |
| *AGAP9* | ENSG00000204172 | 6.27E-19 | 8.56E-14 |
| *AKAP6* | ENSG00000151320 | 2.80E-11 | 4.28E-11 |
| *ANAPC1* | ENSG00000153107 | 9.77E-15 | 2.73E-13 |
| *CD93* | ENSG00000125810 | 1.10E-22 | 6.67E-11 |
| *CEP250* | ENSG00000126001 | 1.85E-11 | 8.29E-12 |
| *CHD3* | ENSG00000170004 | 3.01E-15 | 6.31E-12 |
| *CIT* | ENSG00000122966 | 5.73E-12 | 5.37E-12 |
| *COL6A3* | ENSG00000163359 | 1.15E-12 | 3.55E-11 |
| *CROCC* | ENSG00000058453 | 1.36E-14 | 1.28E-17 |
| *CUBN* | ENSG00000107611 | 1.71E-17 | 2.75E-14 |
| *DNAH11* | ENSG00000105877 | 5.06E-17 | 4.98E-23 |
| *DNAH12* | ENSG00000174844 | 2.28E-12 | 3.08E-15 |
| *DNAH5* | ENSG00000039139 | 1.23E-24 | 3.92E-18 |
| *DNAH6* | ENSG00000115423 | 3.35E-15 | 1.12E-14 |
| *DNAH7* | ENSG00000118997 | 5.30E-17 | 1.25E-19 |
| *DNAH8* | ENSG00000124721 | 1.09E-17 | 2.00E-21 |
| *DNHD1* | ENSG00000179532 | 2.47E-12 | 5.04E-14 |
| *DYNC2H1* | ENSG00000187240 | 1.25E-15 | 9.37E-11 |
| *DYSF* | ENSG00000135636 | 7.77E-13 | 1.53E-15 |
| *ESPN* | ENSG00000187017 | 1.99E-14 | 3.85E-32 |
| *FAT3* | ENSG00000165323 | 1.27E-20 | 2.31E-13 |
| *FAT4* | ENSG00000196159 | 1.27E-14 | 1.69E-14 |
| *FBN2* | ENSG00000138829 | 8.47E-14 | 3.02E-11 |
| *FBRS* | ENSG00000156860 | 7.03E-31 | 6.55E-22 |
| *FRAS1* | ENSG00000138759 | 6.35E-13 | 7.32E-15 |
| *GAPVD1* | ENSG00000165219 | 2.43E-13 | 3.30E-11 |
| *GARRE1* | ENSG00000166398 | 1.50E-15 | 7.96E-14 |
| *HERC1* | ENSG00000103657 | 8.02E-18 | 1.26E-11 |
| *HMCN1* | ENSG00000143341 | 5.70E-22 | 2.27E-24 |
| *IGF2R* | ENSG00000197081 | 1.93E-14 | 1.45E-11 |
| *IQSEC3* | ENSG00000120645 | 4.25E-14 | 2.46E-16 |
| *IRF2BPL* | ENSG00000119669 | 1.33E-30 | 2.67E-11 |
| *ITPR3* | ENSG00000096433 | 1.55E-17 | 4.83E-15 |
| *KIF26A* | ENSG00000066735 | 8.13E-18 | 3.62E-15 |
| *LLGL1* | ENSG00000131899 | 3.10E-12 | 7.71E-11 |
| *LRBA* | ENSG00000198589 | 6.78E-14 | 1.45E-11 |

| | | | |
|---|---|---|---|
| *LRP1* | ENSG00000123384 | 3.01E-19 | 5.04E-15 |
| *MDN1* | ENSG00000112159 | 8.55E-15 | 8.64E-15 |
| *MEGF8* | ENSG00000105429 | 5.47E-15 | 1.44E-11 |
| *MPDZ* | ENSG00000107186 | 8.10E-17 | 1.69E-17 |
| *MYH14* | ENSG00000105357 | 4.64E-13 | 6.59E-15 |
| *MYH6* | ENSG00000197616 | 2.03E-17 | 1.09E-13 |
| *MYH7B* | ENSG00000078814 | 3.06E-15 | 6.69E-14 |
| *MYO15A* | ENSG00000091536 | 1.07E-12 | 1.69E-11 |
| *MYO7A* | ENSG00000137474 | 3.80E-15 | 8.44E-17 |
| *MYOF* | ENSG00000138119 | 7.30E-12 | 8.36E-11 |
| *MYOM2* | ENSG00000036448 | 8.81E-12 | 1.89E-13 |
| *NOTCH2* | ENSG00000134250 | 2.70E-11 | 5.86E-11 |
| *OBSL1* | ENSG00000124006 | 3.41E-19 | 5.77E-14 |
| *OTOF* | ENSG00000115155 | 3.40E-14 | 1.75E-14 |
| *PCDH15* | ENSG00000150275 | 9.69E-14 | 5.58E-12 |
| *PCM1* | ENSG00000078674 | 1.83E-15 | 2.21E-15 |
| *PKHD1L1* | ENSG00000205038 | 3.91E-12 | 2.46E-16 |
| *PPP1R13L* | ENSG00000104881 | 4.25E-14 | 4.13E-11 |
| *RANBP2* | ENSG00000153201 | 9.40E-11 | 9.92E-12 |
| *RGPD4* | ENSG00000196862 | 2.17E-18 | 1.80E-16 |
| *RTEL1* | ENSG00000258366 | 4.08E-17 | 8.42E-15 |
| *RYR2* | ENSG00000198626 | 1.17E-20 | 3.03E-13 |
| *SACS* | ENSG00000151835 | 4.92E-16 | 1.07E-16 |
| *SAMD1* | ENSG00000141858 | 1.08E-11 | 4.66E-13 |
| *SBF1* | ENSG00000100241 | 4.41E-15 | 7.99E-12 |
| *SI* | ENSG00000090402 | 6.32E-20 | 7.60E-11 |
| *SORD* | ENSG00000140263 | 7.26E-11 | 1.11E-11 |
| *SPEG* | ENSG00000072195 | 6.47E-15 | 1.64E-13 |
| *SPTB* | ENSG00000070182 | 7.04E-15 | 3.32E-11 |
| *SZT2* | ENSG00000198198 | 7.96E-18 | 4.90E-14 |
| *UBR4* | ENSG00000127481 | 3.57E-19 | 9.72E-13 |
| *UNC80* | ENSG00000144406 | 6.35E-12 | 8.42E-13 |
| *VPS13B* | ENSG00000132549 | 1.74E-28 | 6.08E-11 |
| *VPS13D* | ENSG00000048707 | 1.96E-22 | 2.40E-15 |
| *VWA8* | ENSG00000102763 | 2.24E-12 | 5.48E-12 |
| *WDR90* | ENSG00000161996 | 2.24E-11 | 4.23E-12 |
| *ZNF462* | ENSG00000148143 | 2.47E-11 | 1.15E-13 |

**Supplemental Figure Legends**

**S1. Pearson correlation between bioinformatic pathogenicity tools and summary variant-level pathogenicity.** Pearson correlation matrix of pathogenicity scores from AlphaMissense, REVEL, ClinPred, PolyPhen, CADD, SIFT, and summary variant-level pathogenicity based on normalised outputs.

**S2. Harmonised scRNAseq UMAP with HLCA level 3 annotation.** Three publicly available single cell RNA sequencing lung datasets representing IPF and control lung samples were harmonised and plotted (Uniform Manifold Approximation and Projection) with annotation at Human Lung Cell Atlas level 3.

**S3. Harmonised scRNAseq UMAP according to disease status.** Harmonised single cell RNA sequencing lung datasets plotted (Uniform Manifold Approximation and Projection) according to disease status of IPF (blue) or control (red).

**S4. Harmonised scRNAseq UMAP according to dataset identifier.** Harmonised single cell RNA sequencing lung datasets plotted (Uniform Manifold Approximation and Projection) according to representative dataset; GSE136831 (blue), GSE135893 (green), GSE128033 (red).

**S5. Flow diagram of burden genes in case cohorts.** Flow diagram summarising the total number of cases within cohorts, and the number of variants represented by overlapping genes according to definition of burden.

**S6. *COL6A3, FAT4* expression by disease status across endothelial, epithelial, immune and stromal lineages.** Plots show p value significance between Control and IPF within Human Lung Cell Atlas level 1 cluster of 20K genes, red IPF, blue non-IPF control.

**S7. *DNAH7, DNAH12, PCM1, MYOF* expression by disease status across endothelial, epithelial, immune and stromal lineages.** Plots show p value significance between Control and IPF within Human Lung Cell Atlas level 1 cluster of 20K genes, red IPF, blue non-IPF control.

**S8. *KIF26A, DYSF, PCDH15* expression by disease status across endothelial, epithelial, immune and stromal lineages.** Plots show p value significance between Control and IPF within Human Lung Cell Atlas level 1 cluster of 20K genes, red IPF, blue non-IPF control.

**S9. *VPS13B, VPS13D, MYOM2, SBF1* expression by disease status across endothelial, epithelial, immune and stromal lineages.** Plots show p value significance between Control and IPF within Human Lung Cell Atlas level 1 cluster of 20K genes, red IPF, blue non-IPF control.

**S10. Relative gene expression in lung across endothelial, epithelial, immune and stromal lineages.** Average and percent expression at Human Lung Cell Atlas level 1.

**S11. Top 10 enriched molecular functions of overlapping protein altering variant genes and exon-level burden genes.** Enrichment analysis for 77 overlapping genes with rare variant burden (A), and 40 overlapping genes with exon-level burden (B). Gene Ontology presents top 10 significant molecular function terms.

**Figure S1. Pearson correlation between bioinformatic pathogenicity tools and summary variant-level pathogenicity.**

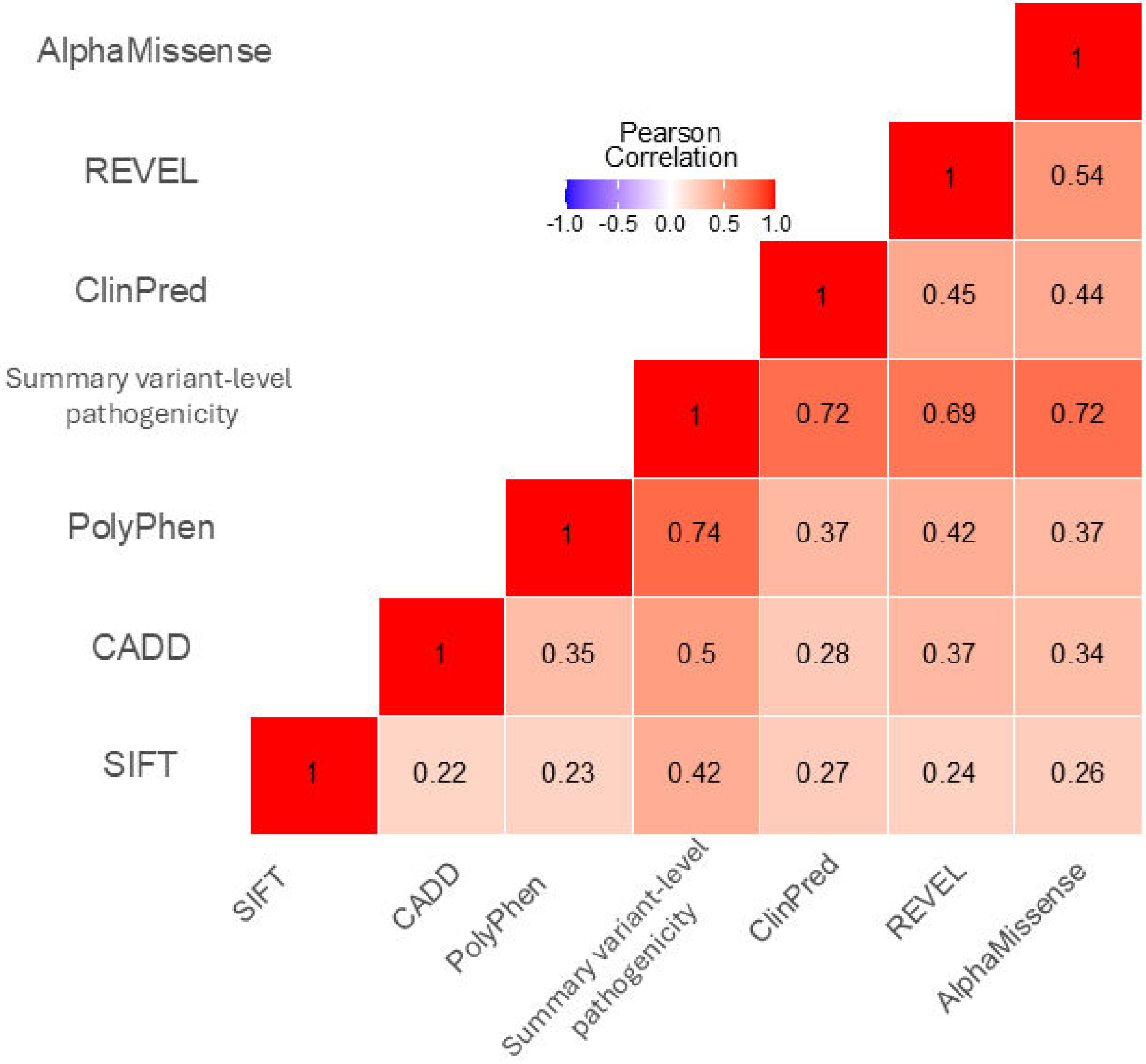

**Figure S2. Harmonised scRNAseq UMAP with HLCA level 3 annotation.**

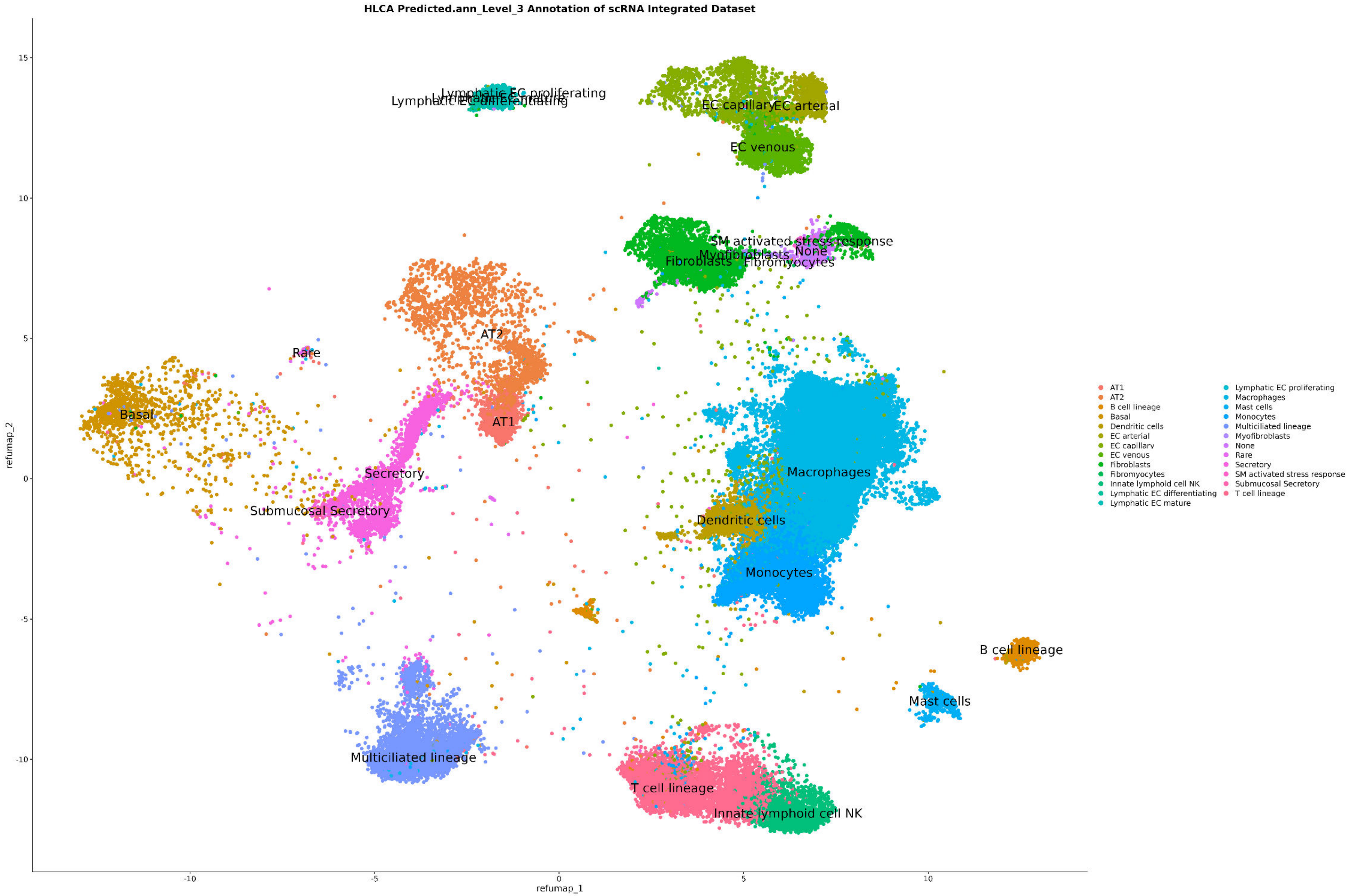

**Figure S3. Harmonised scRNAseq UMAP according to disease status.**

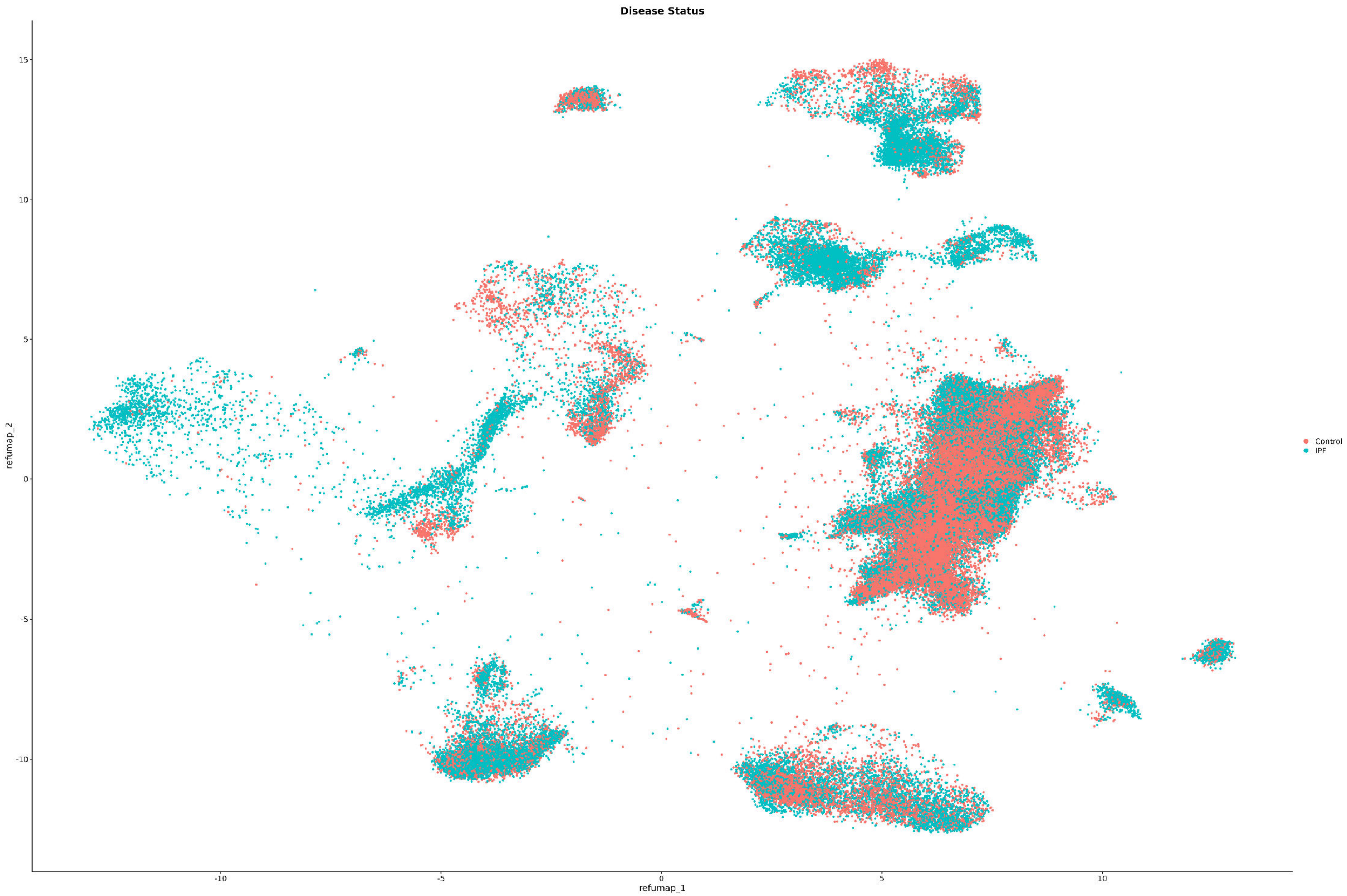

**Figure S4. Harmonised scRNAseq UMAP according to dataset identifier.**

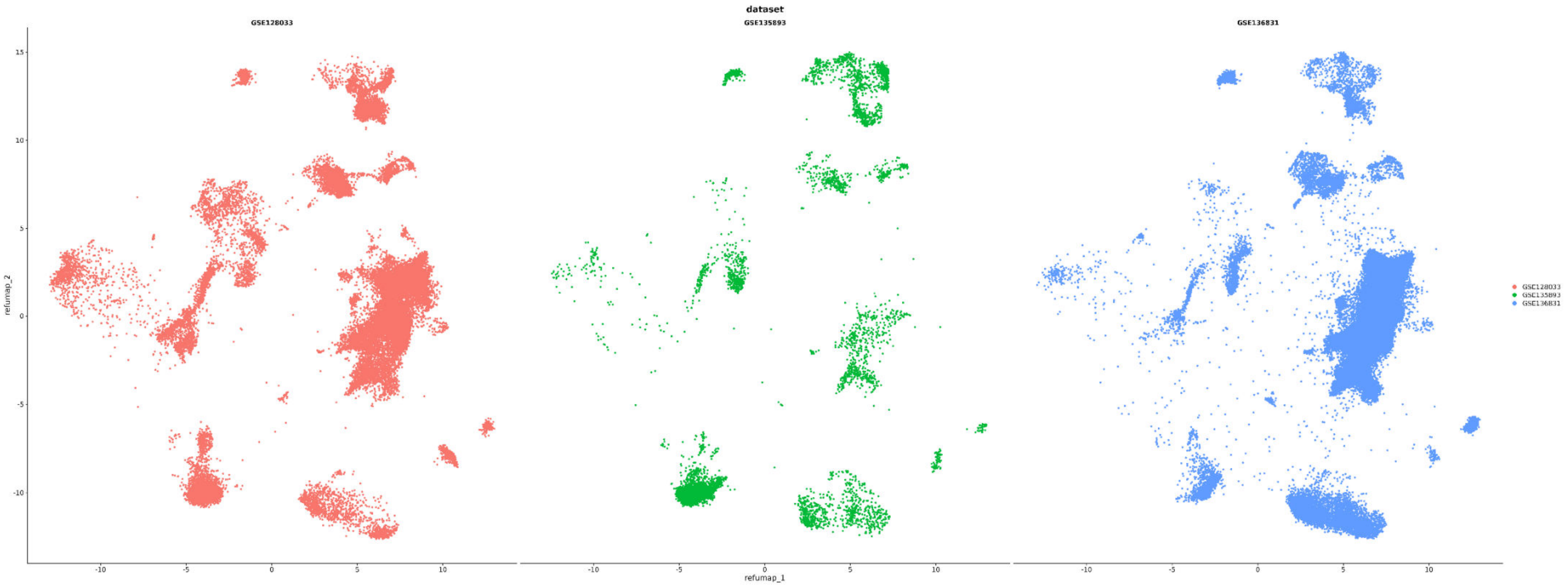

**Figure S5. Flow diagram of burden genes in case cohorts.**

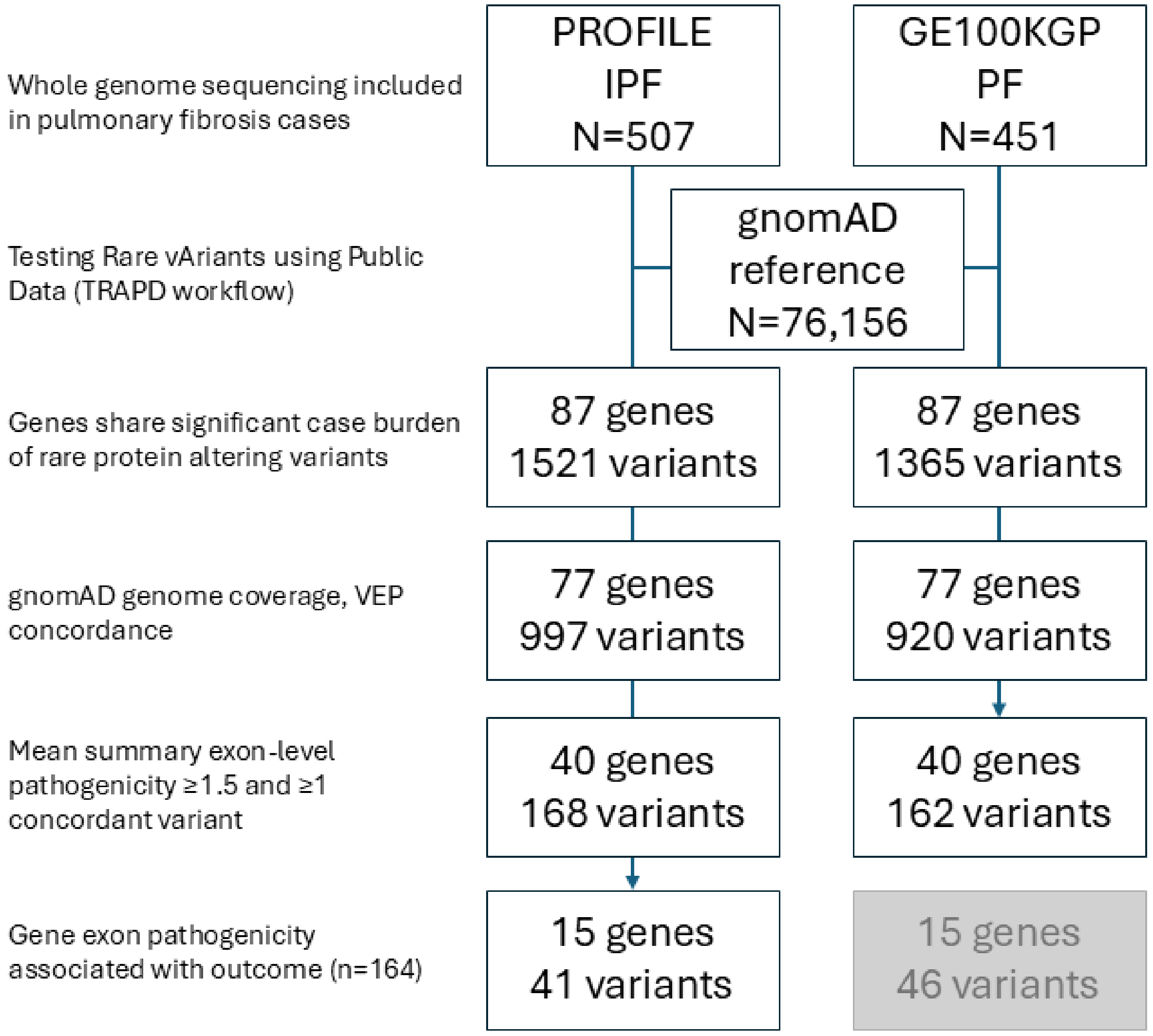

**Figure S6.** ***COL6A3, FAT4*** **expression by disease status across endothelial, epithelial, immune and stromal lineages.**

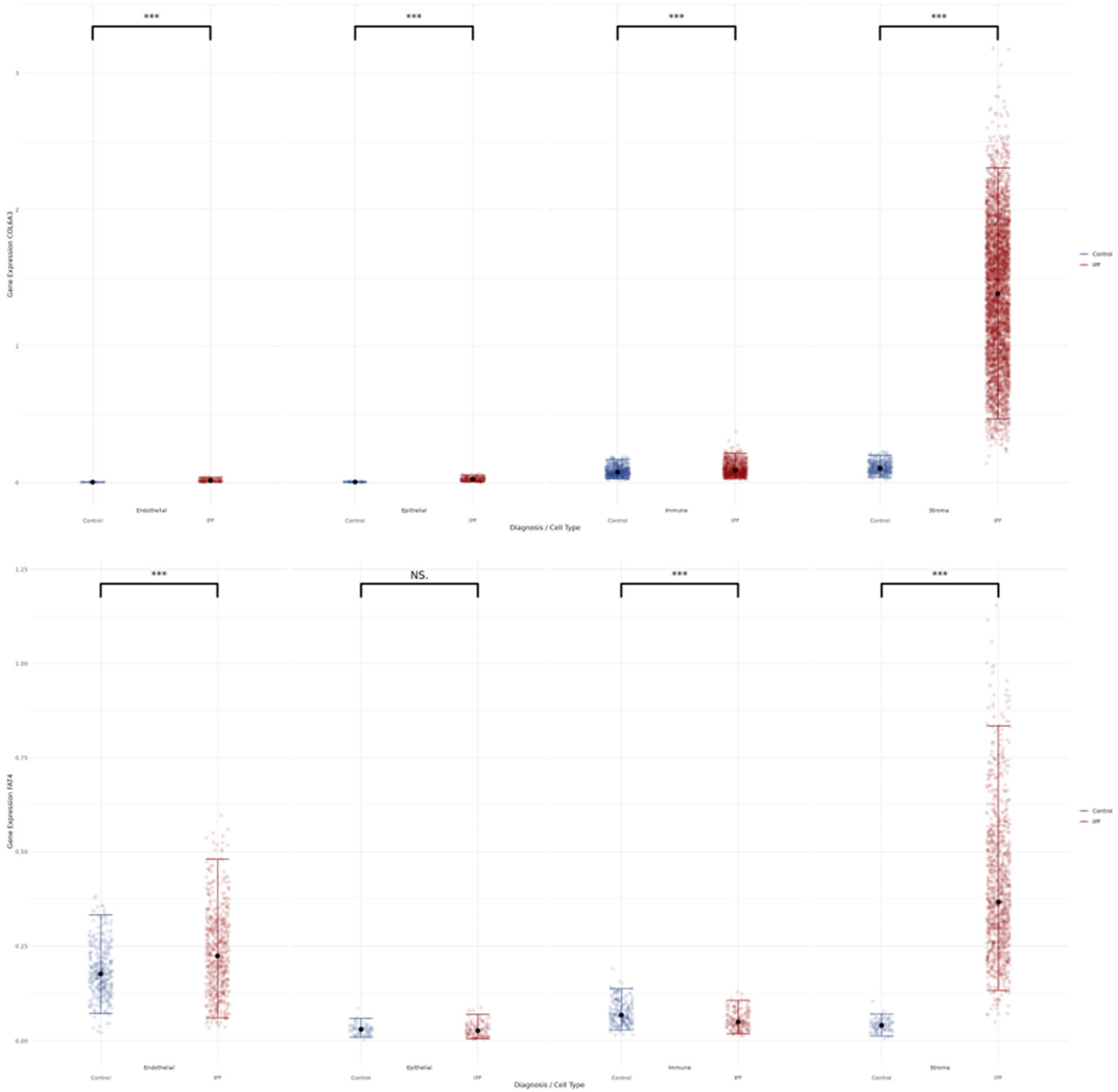

**Figure S7.** ***DNAH7, DNAH12, PCM1, MYOF*** **expression by disease status across endothelial, epithelial, immune and stromal lineages.**

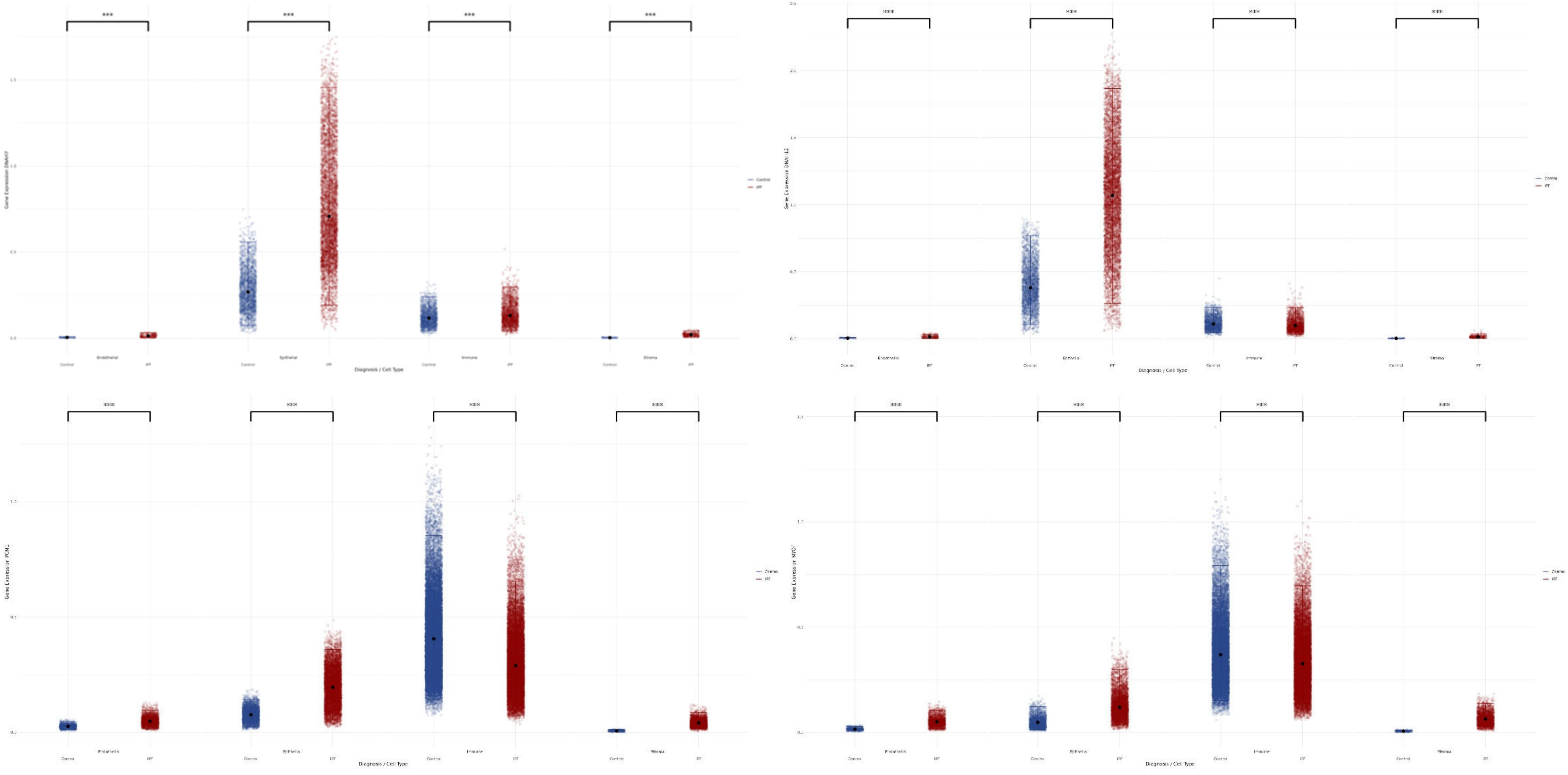

**Figure S8.** ***KIF26A*, *DYSF*, *PCDH15* expression by disease status across endothelial, epithelial, immune and stromal lineages.**

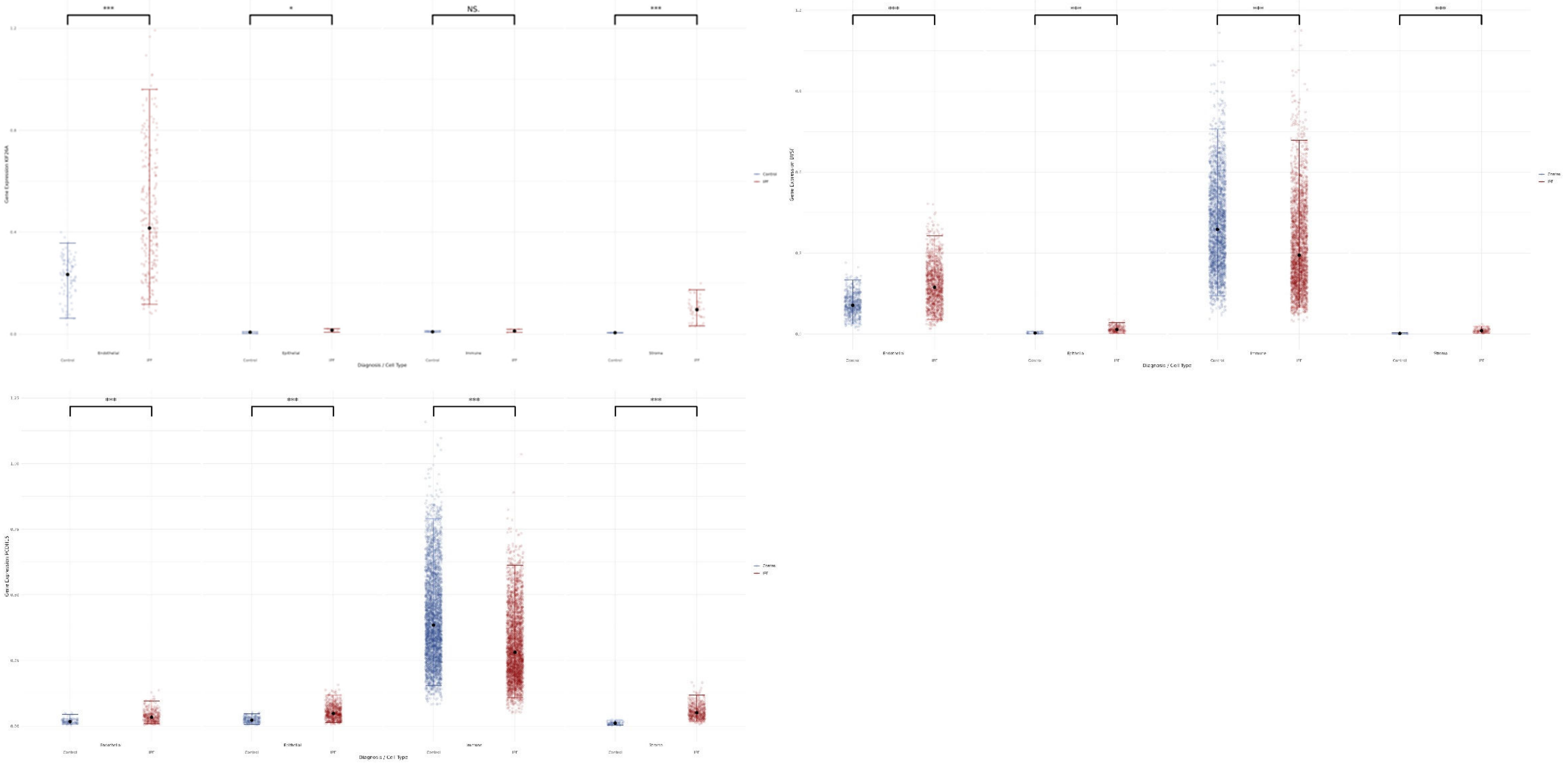

**Figure S9.** ***VPS13B, VPS13D, MYOM2, SBF1*** **expression by disease status across endothelial, epithelial, immune and stromal lineages.**

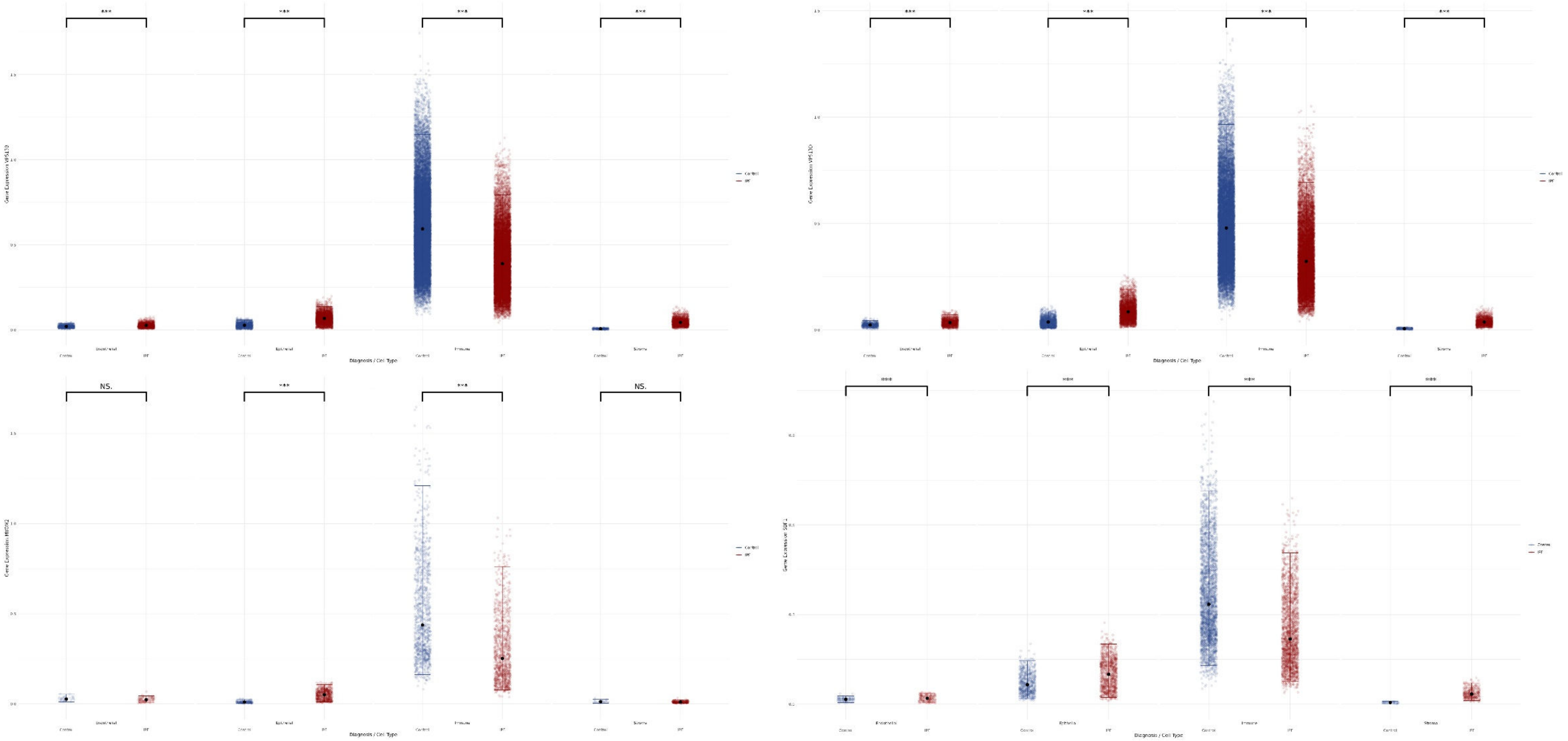

**Figure S10. Relative gene expression in lung across endothelial, epithelial, immune and stromal lineages.**

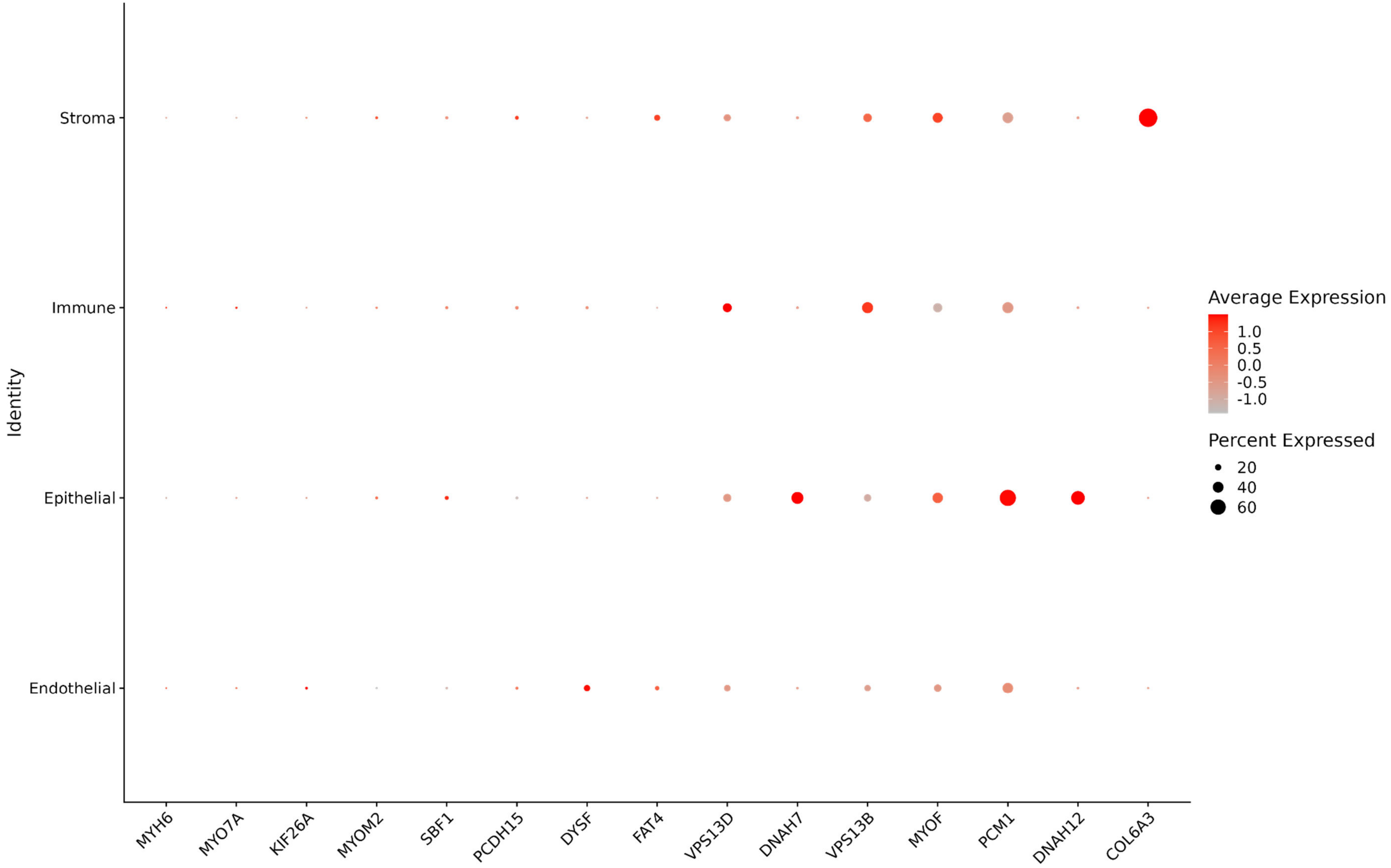

**Figure S11. Top 10 enriched molecular functions of overlapping protein altering variant genes and exon-level burden genes.**

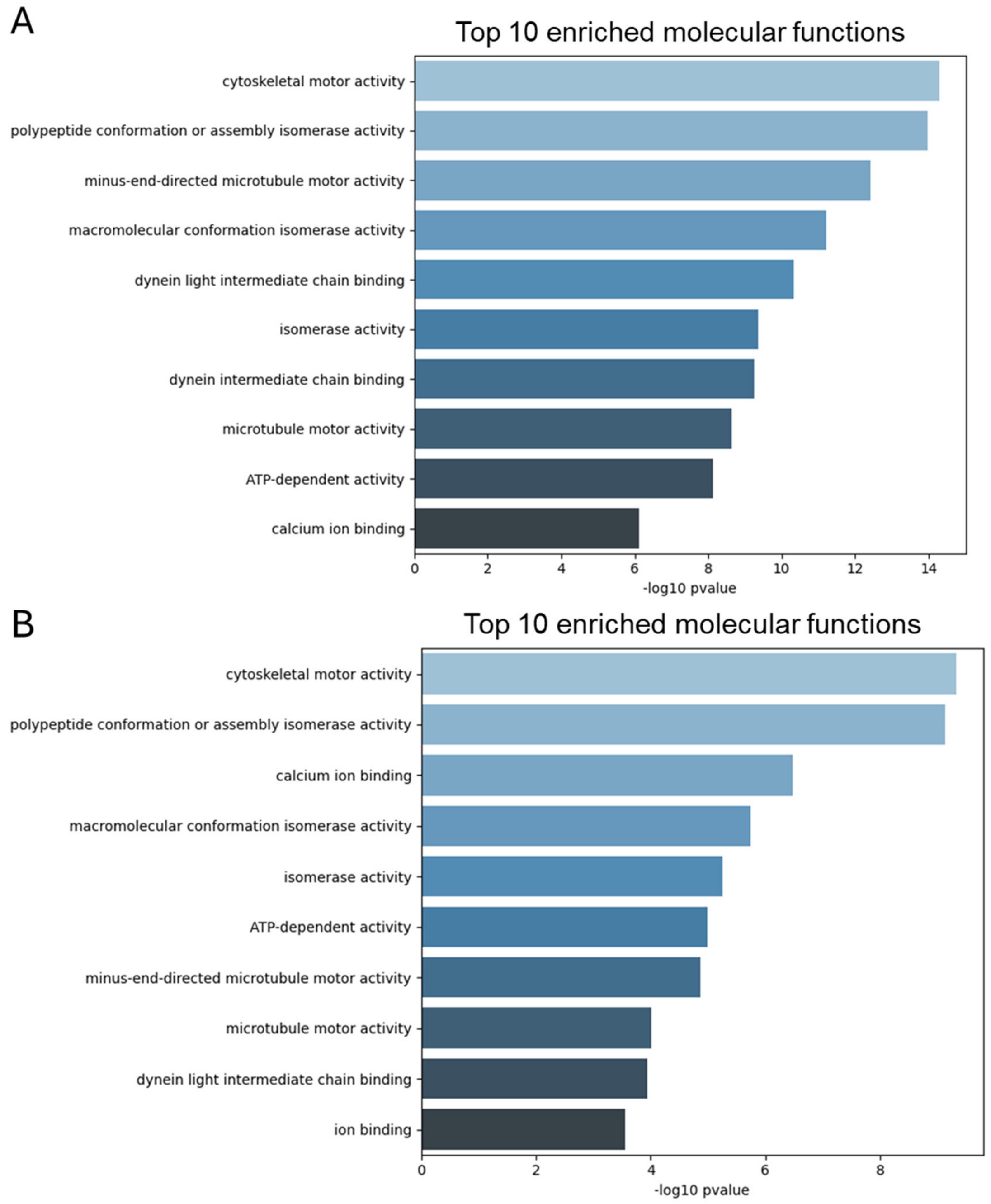
